# Brain-body dynamics are asymmetric and stable across cognitive states

**DOI:** 10.64898/2025.12.22.696055

**Authors:** Jens Madsen, Aimar Silvan, Behtash Babadi, Lucas C. Parra

## Abstract

The human body displays slow, spontaneous fluctuations in brain activity, autonomic physiology, and small incidental movements. It is unknown whether these co-fluctuations reflect a stable endogenous brain-body dynamic, or whether this dynamic varies with cognitive state. We addressed this question using a dynamical systems approach to analyze simultaneously recorded neural activity (EEG), autonomic physiology, and behavior while participants listened to spoken narratives or were at rest. We found that cognitive state did not substantially alter the endogenous dynamic. Acoustic and linguistic features predicted neural activity, which in turn affected physiological responses. Only low-level sound fluctuations exerted direct effects on autonomic signals. Peripheral physiology and behavior exerted stronger influences on EEG than the reverse. These findings suggest that slow co-fluctuations are the result of a stable brain-body dynamic with strong bottom-up feedback, and that the narrative entrains this dynamic by engaging cognition.

## Introduction

The human body is never truly at rest. Even in quiet wakefulness, brain activity, heart rate, pupil size, respiration, and small incidental movements show slow. These fluctuations are often correlated with one another serving well-understood physiological effects. A canonical example is the respiratory sinus arrhythmia (RSA), where heart rate increases during inhalation and decreases during exhalation, reflecting parasympathetic modulation of cardiac function ^1,2^. Pupil size also fluctuates in synchrony with the respiratory cycle, driven by parasympathetic activity and brainstem coordination ^3,4^. These associations have largely been studied in pairs, which neglects the fact that afferent and efferent pathways form a bidirectional network linking the body with a central autonomic network ^5^. How these multiple couplings connect into a coherent whole-body dynamic, and how that dynamic relates to cognition, remains an open question.

Indeed, these physiological signals also correlate with cortical activity, which is more commonly associated with perception, cognition, and behavior. For example, cortical excitability and sensory responses align with the respiratory phase, indicating that respiration entrains neural activity across widespread regions ^6^. Heartbeat-evoked potentials (HEPs) reveal time-locked cortical responses to individual heartbeats, strongest in regions such as the insula and anterior cingulate cortex ^7,8^. Pupil diameter, independent of luminance, correlates with locus coeruleus activity and is often used as an indicator of arousal or mental effort ^9,10^, in particular listening effort ^11^. Increased mental effort has also been linked to increased heart rate ^12,13^. These physiological signals fluctuate at slow frequencies (<0.1Hz), paralleling spontaneous large-scale cortical dynamics measured via fMRI and EEG during rest or unconstrained thought 14,15.

However, these cortical slow fluctuations, which occur even during sleep, cannot be fully attributed to low-level physiology such as fluctuations in CO2 concentrations with breathing and are thought to relate to arousal more broadly ^16^. Together, these observations suggest cortical activity and autonomic physiology form a coordinated dynamical system. Cognitions and emotions are posited to play an important role in these brian-body interactions ^17^. However, we do not know if this dynamic is modulated by cognitive states, or if cognition simply interacts with a stable brain-body dynamic.

Interestingly, auditory perception, in the form of passive listening to narratives, entrains physiological fluctuations including heart rate and pupil size ^18–20^. Continuous natural speech is known to elicit widespread cortical responses, including responses to low-level features such as amplitude envelope ^21^, as well as lexical, semantic, and syntactic information ^22–24^. Narratives therefore provide a natural test case in which cortical processing, autonomic physiology, and behavior are engaged by the same external stimulus. What we do not know is whether auditory narrative listening drives physiological changes, or do arousal-related physiological processes drive cortical activity? Furthermore, does listening impose a new pattern of brain-body interaction, or does it engage with an existing, endogenous dynamic present during rest?

Another factor that has largely been ignored, although it can evidently link arousal to cortical activity, are body movements. The change from rest to active behavior -- the very definition of arousal -- is associated with an increase in heart rate ^1^ and pupil dilation ^25^. Movement will also elicit cortical responses, including incidental behaviors, known to contribute significantly to ongoing brain activity ^26,27^. Eye movements will also elicit cortical activity due to the changing visual input ^28–30^, and will affect pupil size even in the absence of luminance changes ^31^. Is it possible that the association of arousal to cortical activity is due to incidental movement that are often ignored in these studies?

To address these questions, we took here a dynamical systems approach that treats the body and brain as a whole. We modeled interactions of physiological signals with cortical brain activity and incidental movements during processing of an auditory narrative, in an attentive and distracted state, and during silent rest. We hypothesized that co-fluctuations during listening arise from an endogenous dynamic linking cognition with autonomic and oculomotor systems.

To test this, we analyzed multimodal recordings (ECG, respiration, eye movements, electrodermal activity, pupillometry, and EEG, along with acoustic and linguistic features of the narrative. The EEG scalp potentials capture predominantly cortical brain activity ^32^, although we acknowledge subcortical ^33^ and non-neuronal contributions ^34,35^. Focusing on slow fluctuations (down to 0.02Hz), we jointly modeled all signals using a VARX model ^36,37^. This model separates endogenous dynamics from external drive, while allowing for delayed interactions, and extends linear systems models commonly used in neuroscience. ^38–41^ We discover that the endogenous dynamic is largely unaltered by the listening task. The narratives couple into a stable intrinsic brain-body dynamic, with cortical processing mediating the link between stimulus features, physiology, and incidental movements, and these in-turn having strong feedback effects on cortical activity.

## Results

We analyzed neural, physiological and behavioral signals from N=66 healthy adults across two independent experiments in which participants passively listened to emotionally engaging spoken narratives (Fig. 1a). Participants either listened attentively or performed a mental arithmetic task that distracted them from the stories, which continued to play. We also collected data in silence with participants at rest. This design allowed us to examine how slow brain-body dynamics vary between natural listening and resting state, and the effects of attention. Both experiments used the same narrative material and task structure, providing a replication across datasets. They differed only in eye-movement constraints: we either asked participants to look at a central fixation, (Experiment C, ‘constrained’, N=38), or allowed free gaze (Experiment F, ‘free’, N=28). In all instances, the screen was otherwise a blank gray background to minimize visual stimulation. This enabled us to distinguish differences due to gaze behavior from broader physiological influences. We will report here the results for Experiment C and in the supplement for Experiment F. Unless otherwise mentioned, all results replicate across both experiments with identical modeling parameters.

**Figure 1.**
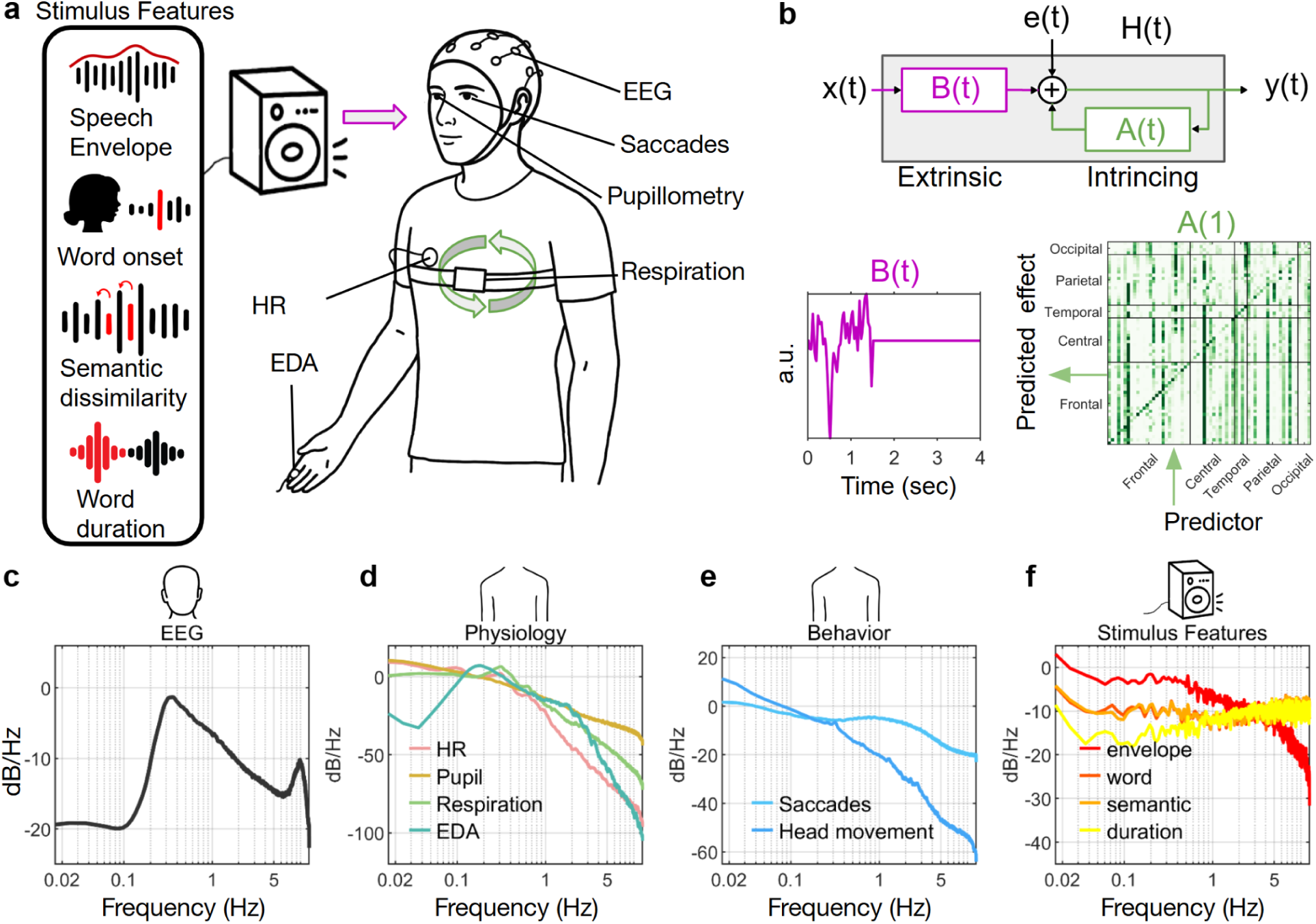
Multimodal recordings and VARX modeling of brain-body dynamics during narrative listening. **a)** Participants listened to naturalistic audiobooks while EEG, pupillometry, saccades, heart rate (HR), electrodermal activity (EDA), and respiration were recorded simultaneously. Acoustic and linguistic features of the narrative were extracted, including the speech amplitude envelope, word onsets, semantic dissimilarity, and word duration. **b)** The vector autoregressive model with exogenous inputs (VARX) separates the overall linear response into extrinsic stimulus-driven filters *B(t)* (purple) and intrinsic recurrent dynamics *A(t)* (green) among all physiological and neural signals. Example filters illustrate the temporal filter of B(t), and the matrix A(t) between EEG channels. **(c-f)** Power spectra of the recorded signals and stimulus features. **(c)** EEG slow-potentials (0.1-5 Hz) show strong infra-slow components dominating the dynamics. **(d)** Physiological signals (HR, pupil diameter, respiration, EDA) also exhibit substantial power at slow timescales. **(e)** Behavioral signals, including saccades and head motion, similarly contain low-frequency structure. **(f)** Speech features span a wide frequency range: the acoustic envelope shows broadband fluctuations, while word onsets, semantic dissimilarity, and word duration include both slow and fast components. The power spectra are consistent across conditions and experiments (Fig. S1).

The multimodal recordings included EEG, heart rate, respiration, pupil size, phasic electrodermal activity, saccade onset times, and head movement power. We focused on raw scalp potentials rather than oscillatory power and refer to these as cortical signals or brain signals and to the other signals, collectively, as body signals. All signals exhibited substantial slow fluctuations in the low-frequency range (0.1-5 Hz, Fig. 1c-e), although we did suppress EEG signals below 0.3Hz for technical reasons (see Methods). This included saccades, despite the instruction to look at the fixation cross in Experiment C (cf. with Experiment F, Fig. S1b). Acoustic and linguistic features of the narratives were extracted to capture both low-level acoustic features, such as the speech envelope and word onsets, and higher-level linguistic information, including semantic dissimilarity and word duration (Fig. 1f).

To quantify how these signals interact, we used a vector-autoregressive model with exogenous inputs (VARX) ^37^. All neural, physiological, and behavioral signals were modeled together as a multivariate time series y(t), while the acoustic and linguistic features of the narrative formed the external input x(t). Filter A(t) describes how signals in y(t) influence one another, whereas the filter B(t) describes the direct impact of x(t) on each modality (Fig. 1b). From these filters we derived effect-size matrices R_A_ and R_B_ that quantify how well each signal predicts another (see Methods). During rest, in the absence of external input, the model reduces to a standard VAR model as used in Granger formalism.^37^ This formulation allows us to quantify endogenous recurrent dynamics and separate that from the coupling with the exogenous narrative drive.

### Endogenous dynamics are unchanged from resting state to listening to a narrative

We first asked whether narrative listening alters the endogenous brain-body interactions that are already present during rest. To test this, we examined the endogenous dynamic captured by the recurrent filters A(t). These filters describe how each neural, physiological, and behavioral signal predicts the others in the absence of external drive. We estimated the effect size matrix R_A_ for brain→brain interactions (Fig. 2a, Fig. S2 for Experiment F), body→body interactions (Fig. 2b), brain-body interactions (Fig. 3, top row), and body-brain interactions (Fig. 3, bottom row, see Fig S3 for Experiment F). If narrative processing imposes new interactions, these matrices should differ between attentive listening and rest. Instead, the endogenous coupling patterns were remarkably similar across states. A permutation test that shuffled R_A_ values showed that the interaction patterns were more similar than expected under random labeling (brain→brain: p < 1·10^−5^, Fig S8a, body→body: p < 1·10^−5^, Fig S8b), indicating that the intrinsic dynamic is not substantially altered by the task.

**Figure 2:**
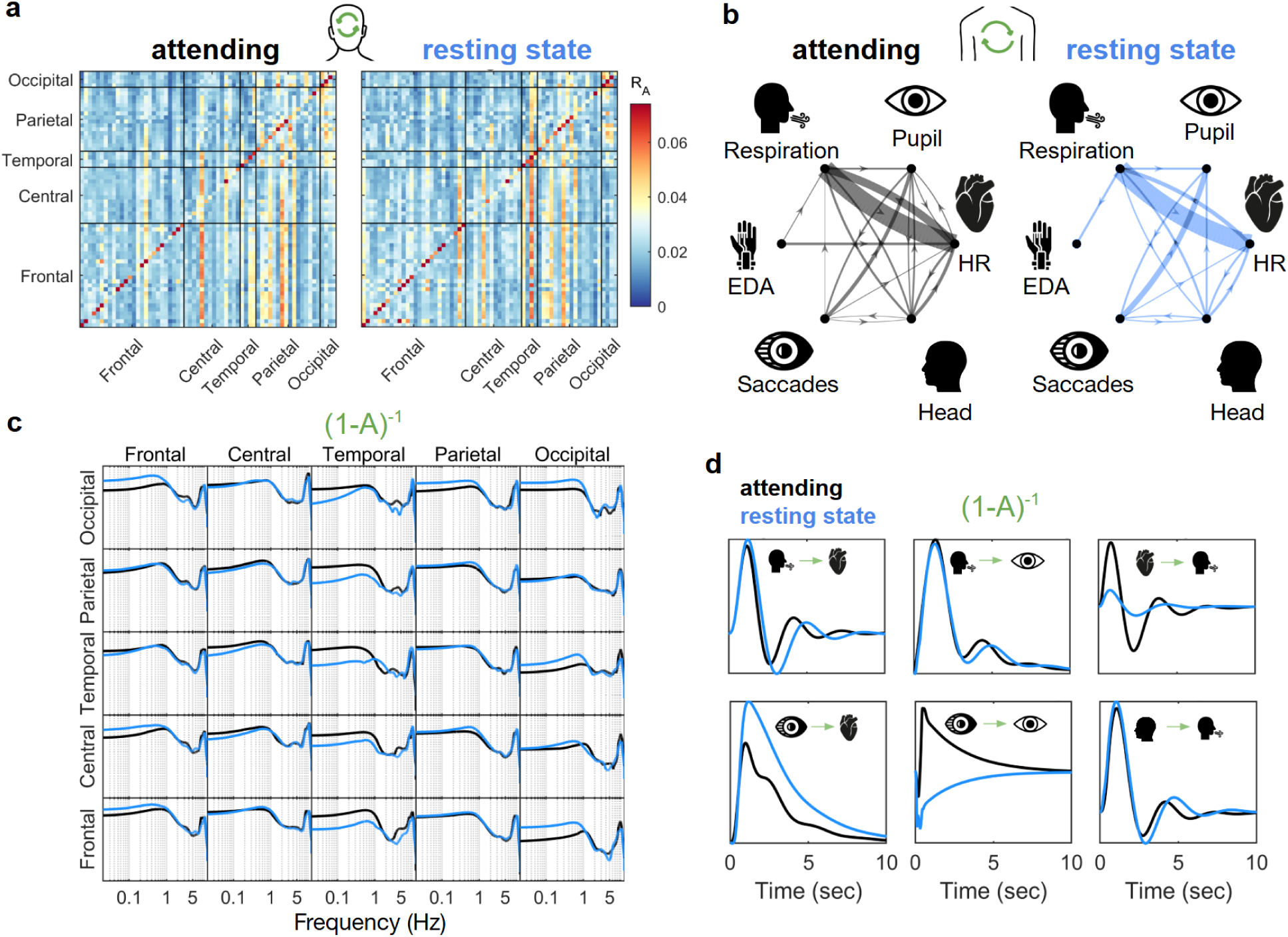
Recurrent dynamics of the brain and the body are largely unaltered by the stimulus vs rest. **(a)** Intrinsic interaction matrices *A* estimated from the VARX model show similar recurrent structure across EEG regions during attentive narrative listening (left) and resting state (right). Each matrix displays the effect size of intrinsic influences among frontal, central, temporal, parietal, and occipital electrode groups. Despite continuous external stimulation during listening, the spatial pattern of endogenous EEG interactions remains highly similar to rest, indicating that slow cortical dynamics are dominated by internal recursion. The horizontal axis indicates the predictors and the vertical axis the predicted effects. **(b)** Intrinsic interactions among peripheral physiological signals also remain stable across conditions. Directed edges represent significant (p<0.001) endogenous influences between respiration, pupil diameter, heart rate (HR), electrodermal activity (EDA), saccades, and head motion. Edge thickness denotes effect size R_A_ (only displaying R_A_>0.02). Both during attentive listening (left) and resting state (right), physiology forms a tightly interconnected network, with respiration and HR showing the strongest endogenous influences and saccades/head motion showing weaker effects. The overall network architecture is preserved across states, demonstrating that narrative engagement does not substantially alter baseline brain-body dynamics. **(c)** Magnitude response from one EEG electrode to another, averaged over all electrode pairs in the corresponding regions (corresponding impulse responses in Fig. S7a). **(d)** Impulse response between body signals. Both (c) and (d) show the recurrent dynamics, symbolically represented as (1-A)^−1^ for the attending and resting state conditions.

**Figure 3.**
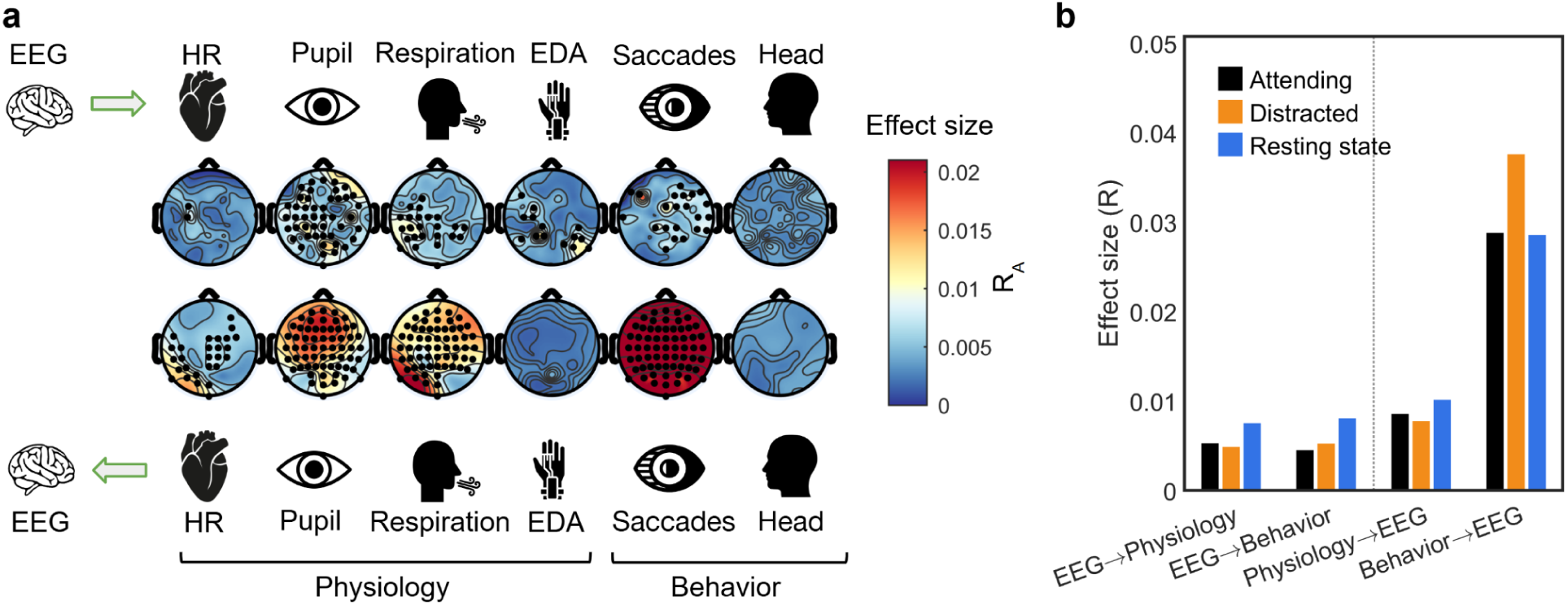
The body affects the brain more than the brain affects the body. **a)** Topographic maps show the effect size of intrinsic interactions between EEG and peripheral signals in the attending condition. The top row depicts the influence of EEG activity on heart rate (HR), pupil diameter, respiration, electrodermal activity (EDA), saccades, and head motion. The bottom row depicts the reverse direction, showing the influence of each physiological or behavioral signal on EEG. Black dots mark electrodes with significant effects. **b)** Grouped bar plots quantify directed effect sizes (generalized R) between EEG and peripheral/behavioral signals across conditions. Values are averaged across electrodes (EEG) and channels within each peripheral/behavioral category. Consistent with panel a, body→brain effects (right group) are generally larger than brain→body effects (left group) across conditions.

Within the brain, R_A_ values showed asymmetric patterns, some electrodes exerted a widespread predictive influence but were themselves weakly predicted (vertical striped pattern in Fig. 2a). Because these patterns depend on sampling rate, reflecting both slow (≤1Hz) and alpha activity (≈10Hz) contributions, we avoid interpreting individual electrodes and instead note the preserved structure of the dynamic (Fig. 2c). This dynamic had two distinctive regimes, one at low frequencies below 1Hz and another at 10 Hz corresponding to the alpha band.

Body-body interactions showed established physiological relationships, validating the approach. Respiration strongly influenced heart rate, which is notably asymmetric, consistent with respiratory sinus arrhythmia (RSA), and saccades influenced pupil size as expected. The model also revealed weaker interactions not widely documented, such as respiration influencing head movements (Fig. 2b). An obvious effect is that of respiration on head movements.

We next examined how perturbations propagate through the endogenous system by computing impulse responses from A(t). These responses unfolded over several seconds and were highly similar between listening and rest (Fig. 2d), including the characteristic asymmetric coupling between respiration and heart rate. The only clear deviation was the saccade-to-pupil effect, which differed between experiments, likely due to differences in gaze behavior under fixation versus free viewing (cf. Fig. 1e and Fig. S1e).

### Stronger bottom-up effects of the body on the brain, than the reverse

We next asked whether cortical activity primarily drives peripheral physiology and behavior, or whether the influence is stronger in the opposite direction. Across all modalities, EEG activity exerted only weak and spatially restricted effects on heart rate, pupil size, respiration, electrodermal activity, saccades, and head movements (Fig. 3a, top row). In contrast, peripheral physiology and behavior produced broad and robust influences on EEG, particularly for pupil diameter and saccades (Fig. 3a, bottom row).

This directional asymmetry was also evident in the overall effect-size (Fig. 3b, see Fig. S3a-b for replication for Experiment F). To determine whether body→brain effects were indeed larger than brain→body effects, we fit the VARX model to each subject and compared the mean effect sizes in matrix R_A_. Body→brain effects were significantly larger across participants than brain→body effects (Wilcoxon’s signed rank test p < 1·10^−6^ for all three conditions; Fig. S8d). While the effect is dominated by saccades driving EEG, the asymmetry remains highly significant after removing saccades from the analysis (Fig. S8e). We conclude that the brain-body dynamic is asymmetric with a dominant effect of physiology and behavior on cortical activity.

We also asked whether the pattern of directed interactions changes with cognitive state. Effect-size matrices for attentive listening and rest were significantly more similar than expected under random relabeling (p < 1·10^−5^, Fig. S8c), indicating that the overall directional structure of brain–body interactions remains stable even when attention varies.

Together, these results show that the dominant direction of influence in the slow brain-body dynamics runs from the body to the brain, and that this directional organization is highly stable across cognitive states. This supports the broader conclusion that narratives couple into an existing endogenous dynamic rather than establishing new pathways, with peripheral physiology and behavior playing a central role in shaping cortical activity.

### Narrative couples with the physiology primarily via cortical activity

We next asked how features of the auditory narrative influence cortical, physiological, and behavioral signals, and whether these effects arise directly from the stimulus or indirectly through the endogenous brain-body dynamic. To quantify this, we examined the extrinsic filter B(t), which captures direct stimulus-driven effects, and compared it to the full impulse response H(t), which includes both direct drive and reverberation through the recurrent intrinsic dynamics A(t) (Fig. 1b). The strength of these direct effects are summarized in the effect size matrix R_B_. The strongest effects of the stimulus are to EEG electrodes (Fig 4a), with the acoustic envelope producing the largest and most spatially extensive responses. Linguistic features, including semantic dissimilarity and word duration, produced weaker effects that were localized to specific scalp regions. By contrast, direct effects on peripheral physiology were very sparse; only fluctuations in speech envelope produced detectable direct effects on heart rate and pupil size (Fig. 4b, purple arrow). All other narrative-related changes in physiology emerged only in the full system response H(t) (Fig. 4e-g), indicating that the stimulus influences peripheral physiology primarily through cortical processing rather than through direct autonomic drive. For example, semantic dissimilarity showed no direct physiological effects in B(t), yet produced clear multi-second responses in heart rate, pupil size, and head motion in H(t), demonstrating that higher-level linguistic information couples to physiology only indirectly via cortical activity, as reflected in the EEG. The few observed direct effects likely involved subcortical nuclei not reflected by the EEG signals, which capture mostly cortical activity. Head movements responded to several stimulus features despite showing no direct stimulus effects and no coupling with EEG (Fig. 3a), suggesting that they arise through indirect pathways mediated by other physiological signals.

**Figure 4:**
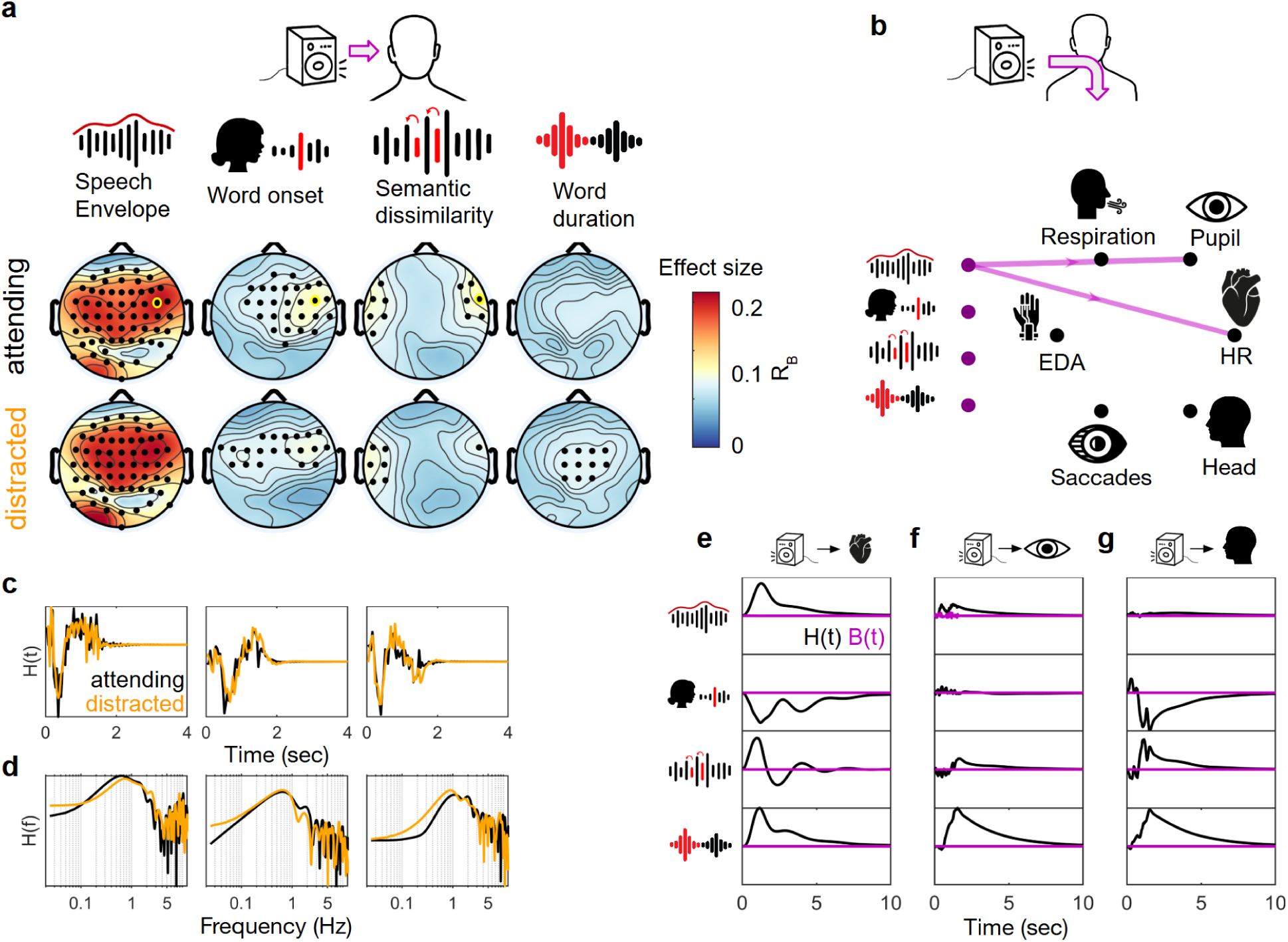
Direct and indirect effects of the stimulus on brain-body dynamics. **a)** Topographic maps show the effect size of stimulus-driven responses from the VARX extrinsic filter B for four speech features: speech envelope, word onsets, semantic dissimilarity, and word duration. Maps are shown for attentive listening (top row) and distracted listening (bottom row). Black dots mark electrodes with significant extrinsic effects. The acoustic envelope produces the largest effects, especially over temporal and fronto-central regions during attentive listening. Linguistic features produce weaker but spatially localized effects. Yellow markers indicate electrode response displayed in c-d. **b)** Network summary of significant extrinsic effects B(t) linking speech features to physiological signals. Lines indicate significant stimulus-to-physiology pathways, with line thickness proportional to effect size. **c)** Time-domain extrinsic impulse responses H(t) for selected significant EEG electrodes (yellow circle in panel a), plotted separately for attentive (black) and distracted (orange) listening. **d)** Corresponding frequency-domain representations H(f), highlighting differences in spectral structure across features and task conditions. **e-f)** Time-domain extrinsic impulse responses B(t) and H(t) for significant (p<0.001) stimulus-to-physiology effects: driving heart rate (**e**), pupil diameter (**f**), and head movements (**g**). All show slow, multi-second H(t) dynamics consistent with autonomic and arousal-linked processes.

Attentive and distracted listening produced highly similar stimulus-locked responses in both amplitude and spectral structure (Fig. 4a, c-d), indicating that the attentional state had only a modest influence on the linear stimulus–response coupling captured by the VARX model. Experiment F showed weaker EEG responses to auditory features than Experiment C (cf. Fig. 4a vs. Fig. S4a), likely reflecting a reduced statistical power due to the smaller sample.

Together, these results show that the narrative engages physiology mainly through its influence on cortical processing, with direct autonomic drive limited to low-level acoustic fluctuations.

### Cortical activity is dominated by intrinsic dynamics with smaller contributions from physiology and behavior

Finally, to compare the relative contributions of stimulus, physiology, behavior, and intrinsic brain dynamics, we quantified how much variance each component explained in the EEG (Fig. 5). External stimulus features accounted for only a very small fraction of EEG variance (approximately 0.1-0.2 percent), with only modest increases during attentive listening. By far the largest contribution (approximately 40-60 percent) came from the EEG itself, reflecting strong spatial correlations across electrodes, likely amplified by volume conduction. Physiological and behavioral signals explained a smaller but reliable fraction of EEG variance (approximately 0.1-1 percent). Among these, behavioral effects were dominated by saccade-related activity, consistent with the large cortical transients produced at saccade offset due to the sudden change in visual input ^42^. Patterns were highly consistent across the two experiments, with the main exception of the behavioral component, which was expected as saccade behavior differed per instructions to the participants (cf. Fig. 5 and Fig. S5).

**Figure 5.**
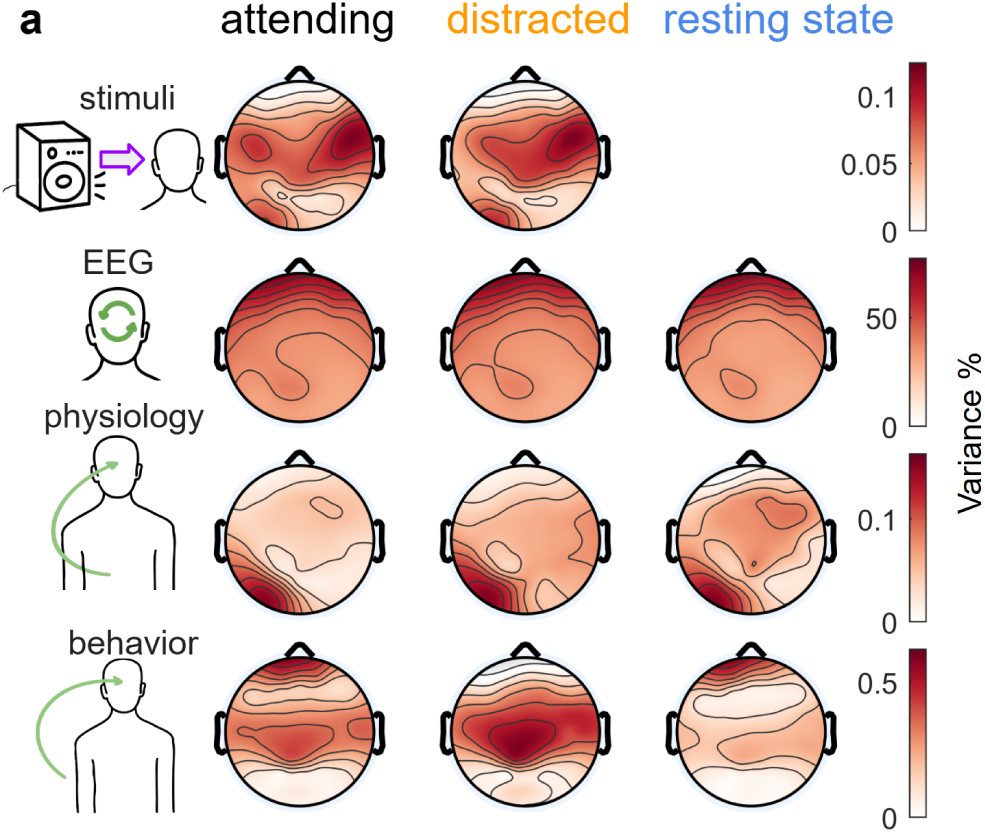
Variance of the EEG signal is dominated by the brain, and to a lesser degree by behavior and physiology, with only minor effects from the stimulus. Topographic maps show the percentage of EEG variance explained by four sources in the VARX model: external stimulus features (top row), intrinsic EEG dynamics (second row), intrinsic influences from peripheral physiology (third row), and behavior (bottom row). Maps are shown for attentive listening, distracted listening, and resting state.

Together, these results show that cortical activity during naturalistic listening is shaped overwhelmingly by internal brain dynamics, with smaller contributions from oculomotor and other peripheral signals, and only minimal direct influence of the external stimulus.

## Discussion

We set out to understand how slow, correlated fluctuations in neural, autonomic, and behavioral signals relate to cognition and external stimulation. A central question was whether these fluctuations reflect a stable endogenous dynamic or whether they reorganize when cognitive processes such as narrative comprehension are engaged. We found that listening to auditory narratives did not substantially alter the recurrent interactions observed during rest. Instead, narrative features are coupled into the same intrinsic structure present in the resting state, indicating that the overall organization of brain-body interactions remains stable across cognitive conditions. This directly addresses the question of whether slow fluctuations reconfigure with task demands and suggests that they do not.

A second question concerned how narrative features influence physiology. Although several acoustic and linguistic properties of the narrative evoked measurable physiological responses, the pattern of effects revealed a clear hierarchy. Low-level acoustic fluctuations produced the only direct autonomic responses, whereas higher-level linguistic features influenced physiology only indirectly through cortical activity. This distinction aligns with known neurophysiology: sound-level fluctuations may access autonomic pathways through brainstem nuclei ^9^, while semantic signals appear to require cortical mechanisms involved in speech comprehension, such as those described by ^22,23^. In this view, cognitive-level processes shape physiology predominantly via cortical processing, while autonomic pathways can respond to basic acoustic energy even without cortical involvement.

At the same time, the influence of physiology on cortical activity was substantial. Across modalities, peripheral physiology exerted a stronger influence on EEG than the reverse.

Pupil-linked arousal predicted fronto-central EEG activity, consistent with prior work on noradrenergic modulation of cortical excitability ^9,43^. Respiration broadly influenced slow cortical fluctuations, in line with fMRI evidence for respiration-linked global signals ^44,45^. Heart-rate fluctuations affected posterior EEG channels in a manner consistent with heartbeat-evoked responses ^46,47^. Together with recent fMRI findings showing widespread cortical modulation by autonomic activity ^16^, these results suggest that slow neural fluctuations may reflect an endogenous brain-body dynamic.

A related question was whether previously reported associations between arousal and cortical activity might be partly attributable to incidental movements. Despite instructions to maintain fixation, participants nonetheless moved their eyes almost as much as during the unconstrained condition, perhaps “carried away” by the primary task of listening to the story. Those eye movements, over a blank gray screen, still exerted a substantial influence on the EEG. This is consistent with widespread neural responses following fixation onset ^28–30,48^. Head movements also responded robustly to narrative features, yet showed no direct coupling to EEG or the stimulus; instead, they appeared to reflect influences from other physiological signals. These findings highlight an important consideration for interpreting arousal effects: multimodal measurement is essential to distinguish true arousal-related cortical modulation from sensory consequences of movement associated with physiological arousal.

In addition to these movement-related effects, cortical activity itself was dominated by slow potentials at or below 1 Hz, which showed strong spatial correlations across the scalp. These slow components likely reflect a mixture of volume conduction and genuine large-scale distributed neural processes ^49^. The persistence of these slow fluctuations across resting, attentive, and distracted listening reinforces the idea that intrinsic dynamics set the dominant timescale of brain activity, with stimulus-driven influences adding only modest contributions.

Because intrinsic dynamics dominate slow cortical activity, narrative-driven effects on physiology and behavior must be interpreted within this pre-existing structure. Head movements, for example, showed no direct coupling to EEG or the stimulus, yet responded robustly to multiple narrative features. This indicates that head motion was influenced through its connections with other physiological signals that were themselves modulated indirectly by the narrative. These patterns highlight that naturalistic stimuli do not propagate through isolated feedforward pathways; rather, narrative features enter an already structured brain–body dynamic in which cortical, autonomic, and behavioral processes interact continuously.

Although intrinsic cortical dynamics account for most of the variance in the EEG, the directed influences we observed were asymmetric: peripheral physiology and behavior predicted EEG activity to a higher degree than the reverse. This indicates that the slow cortical fluctuations are substantially driven by autonomic and oculomotor activity. This interpretation is consistent with theoretical views emphasizing the central role of interoception and autonomic regulation in shaping cortical processing^50^ and with empirical observations on the effect of autonomic signals on cortical activity^12,16,45^ and cognition ^51^ In this view, bodily signals form part of the endogenous context within which cortical dynamics evolve.

Taken together, our results indicate that narratives influence physiology by recruiting an endogenous dynamical system, and not via any unique regulatory processes. This extends prior findings that endogenous cortical dynamic remains largely unchanged between rest and watching movies ^36^ or tasks states ^52^ and supports theoretical accounts emphasizing the primacy of internal dynamics in neural processing ^53^. Our analysis of brain-body interactions as a dynamical system is in line with recent emphasis in neuroscience that regards the brain as an adaptive dynamical system. ^54,55^

We set out to understand the source of slow coordinated fluctuations observed in physiology and neural signals. We modeled interactions as a recurrent system with delays and found strong feedback effects. An established result in dynamical systems theory is that a strong feedback with delays in a bounded (nonlinear) system will tend to fluctuate, or even oscillate^56^. The fluctuations in HR at a resonance frequency of 0.1Hz have been ascribed to such dynamic effects^57,58^ . As the bottom-up feedback is strongest in the low frequencies, we argue that a similar mechanism can be the source of the slow fluctuations observed in body and brain more rapidly.

In our view, the purpose of the endogenous brain-body dynamical system is to support allostasis^59^. The notion of allostasis is that the brain predicts the demands of the body and drives physiology accordingly in preparation. This is perhaps best researched in the context of heart rate fluctuations. For instance, heart rate decelerates when we need to orient ourselves to salient external events, or when we need to prepare to act or adjust our behaviour ^60^. Heart accelerates when imagining movement, in proportion to the imagined effort of that movement ^61–63^. Mental imagery, by relieving arousing emotions like fear or anger, also increases cardiac activity ^64–66^, to a bigger extent than sad, relaxing, or pleasant emotion imagery ^67^. Perhaps narratives similarly drive these autonomic responses by driving our imagination. In this alternative point of view, the slow fluctuations observed during quiet rest may reflect mind wandering ^68^. Spontaneous thoughts may engage the same neural-autonomic circuitry as externally driven narrative imagery with strong bottom-up feedback. Understanding how this intrinsic dynamic shapes perception, cognition, and bodily regulation may offer a unifying framework for interpreting variability in neural and physiological signals across conditions and individuals.

### Contrast with existing literature and limitations of this study

A few results may have been at variance with existing literature, but we ascribe these differences mostly to methodological differences. For instance, a number of studies have analyzed EEG activity together with heart rate and respiration using various pairwise correlation measures ^69–72^. Consistent with our results, these studies find extensive links between EEG and physiological signals, and in particular a strong effect of heart-rate variability on EEG ^71^, but the connectivity seems to be more variable than what we have found ^69,72^. These studies differ from the current work in that they analyzed oscillatory band-pass power of the EEG, rather than the slow potential fluctuations as we have done here.

We did not observe substantial differences between attentive and distracted conditions using the linear VARX model. Whereas prior work using inter-subject correlation has shown stronger coupling of the brain and physiology with the narrative activity during attentive listening ^18,73–80^. The discrepancy likely reflects methodological differences. Inter-subject correlation captures any shared neural response, including nonlinear responses with long temporal dependencies, whereas the VARX model assumes linear relationships with relatively short temporal windows (here in the order of seconds). Cognitive processes such as narrative comprehension may involve nonlinear integrations over longer timescales of minutes ^23,81^, which are not captured with the present model parameters.

Several methodological and interpretive considerations should be taken into account when evaluating these findings. First, although we interpret the EEG signal as reflecting predominantly cortical activity, slow components of the EEG can include non-cortical and non-neuronal contributions. The endogenous interactions we observe extend to frequencies near and below the 0.3 Hz high-pass cut-off used for artifact handling. Even though these components have been filtered, residual activity is still present and may reverberate within the intrinsic dynamics. Slow voltage fluctuations can arise from galvanic skin potentials or respiration-related artifacts ^82^, as well as from established slow cortical potentials such as the contingent negative variation ^82,83^.

Second, several results depended on modeling parameters. Reducing the sampling rate from 25 Hz to 10 Hz removed alpha-band activity and substantially reduced effect sizes R_A_ between electrodes, even though the overall variance explained by other electrodes changed little. This suggests that low-frequency EEG drives most of the predictive power, whereas the detailed spatial coupling patterns depend on oscillatory structure such as alpha coherence. Model order selection using the Akaike information criterion yielded a clear optimum for the exogenous filter order n_b_, but not for the autoregressive order n_a_, which decreased approximately linearly with larger values. This pattern likely reflects nonlinear long-term memory effects that are not well captured by linear autoregression.

Third, some findings depended on choices in behavioral preprocessing. Coding saccades by onset alone emphasized their motor component, whereas including saccade length produced stronger occipital responses consistent with fixation-onset visual input ^30,84^. Differences in saccade-to-pupil impulse responses between experiments also suggest sensitivity to measurement choices, and detailed time courses should therefore be interpreted with caution.

Fourth, heart-rate effects on posterior EEG channels may reflect heartbeat-evoked responses ^46,47^ or vagal pathways, but resolving these mechanisms would require methods sensitive to subcortical sources and non-linearities.

Finally, linear models are used extensively in neuroscience, including recurrent ^41^, feed-forward ^40^, and first order dynamical models.^38^ The VARX model extends over these transitional approaches by allowing for delayed interactions (i.e. n_a_>1), or by combining recurrent with feedforward dynamics. However, while one can argue that linear models suffice to capture large scale dynamics^39^ we acknowledge that it cannot capture nonlinear effects (boundedness) or long-context effects known to be important in cognition. Future work incorporating nonlinear or state-space models may better capture these extended temporal dependencies and their relationship to autonomic and cortical dynamics ^16,44,45^.

## Author Contributions

J.M. and L.C.P. conceived and designed the study. J.M. collected the data. J.M. and A.S. analyzed the data. B.B. contributed to the development of statistical analyses. J.M. and L.C.P. wrote the manuscript, with input from all authors.

## Acknowledgments

The authors acknowledge support from the National Science Foundation through grant no. DRL-2201835 and funding from the National Institute of Health grant no. P50 MH109429.

## Methods

Resource Availability

### Lead Contact

Further information and requests for resources, data, or analysis code should be directed to the lead contact, Dr. Jens Madsen.

### Materials Availability

No new biological reagents were generated for this study.

### Data Availability

All neural and physiological recordings used in this study were collected previously as part of an earlier experiment, but the full dataset has not been publicly released. The original published work ^19^ reported analyses of EEG, eye-tracking, ECG, and respiration. Electrodermal activity (EDA) was recorded during the same experimental sessions but was not analyzed or reported previously. The same is true for various features extracted from the auditory narrative. Data and analysis code will be made available upon publication.

### Participants

Data were collected as part of two experiments conducted at the City College of New York. Experiment C included 43 participants (31 female, age range 18–30 years, M = 21.58, SD = 2.81), of whom 5 were excluded due to poor signal quality or timing anomalies. Experiment F included 32 participants (16 female, age range 19–36 years, M = 23.69, SD = 4.42), with 4 exclusions for similar reasons.

### Ethics statement

All procedures were approved by the Institutional Review Board of the City University of New York. Informed consent was obtained from all participants prior to participation.

### Stimuli and experimental procedure

Participants listened to a curated set of ten emotionally engaging autobiographical audio narratives, originally used by ^85^, and selected from public archives including StoryCorps and The New York Times *Modern Love* series. Each story ranged from 2 to 5 minutes, totaling 27 minutes of narrative content.

In the resting state condition, participants sat with eyes open, viewing either a blank gray screen (Experiment F) or a gray screen with an isoluminant fixation cross (Experiment C), to maintain comparability with the auditory-narrative conditions.

Participants were seated comfortably in a sound-attenuated booth with white cloth walls and ambient LED lighting. Auditory stimuli were delivered via stereo speakers positioned at approximately ±60° azimuth relative to a 27-inch monitor placed ∼60 cm away. In Experiment C, a central isoluminant fixation cross was displayed to encourage ocular stability, whereas Experiment F used a blank gray screen.

The experiment included two listening conditions. In the attentive condition, participants were instructed to listen carefully to the narratives while maintaining visual fixation. In the distracted condition, participants silently counted backward in steps of seven from a random prime number between 800 and 1000, ensuring that the auditory stimulus remained identical while cognitive engagement differed.

Across all participants, this resulted in the following total recording durations:

- Experiment F (N = 28): 752.40 minutes of attentive and distracted listening, and 139.98 minutes of resting state.
- Experiment C (N = 38): 1,021.11 minutes of attentive and distracted listening, and

190.03 minutes of resting state.

### Signal acquisition and synchronization

To ensure multimodal alignment, auditory onset and offset triggers were embedded in all stimulus files and logged via a Cedrus StimTracker, including an audible beep at the start and end of each story. All modalities were timestamped using the Lab Streaming Layer (LSL) protocol. Additional hardware triggers were distributed to both EEG and eye-tracking systems. Linear regression models were used to temporally align data streams post hoc using these shared triggers.

### Eye-tracking and pupillometry

Eye position and pupil diameter were recorded using the SR Research EyeLink 1000 system with a 35-mm lens at a sampling rate of 500 Hz. A standard 9-point calibration was followed by manual verification. To avoid visual artifacts, all visual stimuli were adjusted to be isoluminant. Head movement was unconstrained, allowing for natural posture. Saccades and blinks were detected using the device’s built-in algorithm. Blinks and saccades, along with 100 ms of pre-and post-event data, were linearly interpolated. Pupil diameter signals were then downsampled to 100 Hz. Temporal Response Functions (TRFs) were estimated for blink and saccade offsets (−4 to +5 s), and their linear contributions were regressed out. Segments that had more than 50% data missing were removed and not included in the analysis. Gaze position, gaze variance, and pupil size signals were subsequently downsampled to 25Hz. The saccade signal was coded as a single-sample spike train (0/1) signal at the saccade onset.

### EEG recording and preprocessing

EEG was recorded at 2048 Hz using a BioSemi ActiveTwo system with 64 scalp electrodes placed according to the 10-10 international system. Four electrooculogram (EOG) electrodes were placed above, below, and lateral to the eyes to capture ocular artifacts. Robust Principal Component Analysis (Robust PCA; ^86^) was applied for artifact removal in both EEG and EOG channels.

EEG data were band-pass filtered (0.3-64 Hz), downsampled to 25 Hz, and manually inspected for noisy channels. Interpolation of bad channels was conducted using neighboring electrodes in a 3D-projected coordinate space. Eye-movement artifacts were further removed via linear regression of EOG channels from EEG data. Cardiac artifacts were removed using regression of ECG-derived regressors. Residual outliers exceeding ±4 interquartile ranges were replaced by interpolation using neighboring electrodes ±40 ms. The 0.3 Hz high-pass filter was necessary for spatial interpolation and artifact rejection procedures to operate effectively. However, interactions among electrodes extended into frequencies below this cutoff. Power below 0.3 Hz was attenuated by approximately 20 dB relative to activity at 1 Hz (corresponding to ∼10× smaller amplitude), but these slow components were still present and contributed to the recurrent dynamics modeled in the VARX analysis. Such infra-slow contributions may include slow cortical potentials as well as non-neuronal sources such as galvanic skin responses or respiration-related shifts ^34,35^.

### ECG recording and preprocessing

ECG was recorded using the BioSemi ActiveTwo system at a sampling frequency of 2048 Hz. Two electrodes were placed below the left clavicle and one on the left lower torso, forming a standard Lead II - like configuration. The raw ECG signal was high-pass filtered at 0.5 Hz to remove slow drifts and notch filtered at 60 Hz to suppress line noise. R-peaks were detected using the *findpeaks* function in MATLAB (MathWorks), with peak-prominence and minimum-distance parameters adjusted per subject to ensure robust detection. Instantaneous heart rate (HR) was computed as the inverse of the inter-beat interval for each pair of successive R-peaks. The resulting irregularly sampled HR time series was then interpolated onto a uniform sampling grid using cubic spline interpolation. For consistency with all other physiological and neural signals used in the VARX model, the HR signal was resampled to 25 Hz. All preprocessing steps were performed separately for each participant prior to statistical analysis.

### Electrodermal activity (EDA) recording and preprocessing

Electrodermal activity was recorded using the BioSemi ActiveTwo GSR sensor module, which employs a pair of non-polarizable, pure sintered Ag/AgCl electrodes. Electrodes were affixed to the index and middle fingers of the non-dominant hand using adhesive rings and isotonic electrode gel, following manufacturer recommendations to avoid alcohol-based skin preparation. The signal was digitized at 2,048 Hz through the ActiveTwo auxiliary input.

EDA signals were first low-pass filtered at 10 Hz to remove high-frequency artifacts. We extracted the phasic component of the EDA signal, which reflects rapid sympathetic responses, by applying a two-stage band-pass filter: a 5th-order high-pass filter at 0.16 Hz to remove tonic drift, followed by a 5th-order low-pass filter at 2.1 Hz. Filtering was performed using zero-phase IIR filters implemented in second-order sections. The phasic component was then downsampled to 25 Hz to match the sampling rate of the other physiological and neural variables used in the VARX model.

### Respiration recording and preprocessing

Respiratory effort was measured using a piezo-crystal respiration belt (SleepSense 1387; BioSemi ActiveTwo), placed around the chest to capture changes in thoracic circumference. The signal was digitized at 2,048 Hz through the ActiveTwo auxiliary input. Because the piezo element produces a polarity that depends on belt orientation, signal polarity was manually corrected when necessary to ensure that inhalation corresponded to positive deflection.

No additional preprocessing was applied. The raw respiration waveform was downsampled to 25 Hz to match the sampling rate of the other physiological and neural signals included in the VARX model. For visualization purposes, we also computed an analytic envelope of the Hilbert transform to illustrate respiratory depth over time, but statistical analyses used the downsampled raw signal.

### Head-movement recording and preprocessing

Head position was recorded continuously using the SR Research EyeLink 1000 system, which tracks head position with a camera at a sampling rate of 500 Hz. Participants were seated without a chin rest to allow natural posture and small incidental movements. Raw head-position signals were exported in three spatial dimensions (x, y, z). To remove slow postural shifts, each dimension was high-pass filtered at 0.1 Hz using a 5th-order Butterworth filter. Instantaneous head-movement magnitude was then computed as the root-mean-square amplitude of the analytic signal obtained via the Hilbert transform, with mean taken across the three spatial dimensions. The resulting time series was downsampled to 25 Hz to match the sampling rate of the other physiological and neural signals used in the VARX model.

### Speech feature extraction

Four speech-derived regressors were computed to characterize acoustic and linguistic drive. *Acoustic envelope* was obtained by computing the Hilbert amplitude of the audio waveform and downsampling to 25 Hz.

*Word onsets* were coded as a pulse train marking the onset of each word in the transcript. *Word duration* was represented as a single pulse at the onset time with amplitude equal to the upcoming word’s duration.

*Semantic dissimilarity* was computed following the framework of Broderick et al. (2018), using MATLAB’s fastText word embedding model (*fastTextWordEmbedding*). For each sentence, words were embedded using *word2vec*, and semantic dissimilarity was defined as one minus the correlation between the current word embedding and the mean embedding of the preceding words in that sentence. For the first word of each sentence, the mean embedding from the previous sentence was used; if no prior sentence existed, the value was left undefined. These dissimilarity values were time-aligned to word onsets and downsampled to 25 Hz for use as exogenous regressors in the VARX model.

All speech features were temporally aligned with EEG and physiological recordings at 25 Hz.

### Quantification and Statistical Analysis

#### Dynamical modeling with vector-autoregressive models with external input (VARX)

The approach we used here is similar to the temporal-response function approach often used for modeling responses to continuous speech ^21,40^. It is similar in that the model captures linear impulse responses H(t) to the stimulus. It does differ in that the linear multi-input-multi-output impulse response H(t) is uniquely factored into a moving average filter matrix B(t) and autoregressive filter matrix A(t) (Fig. 1D) ^36,37^.

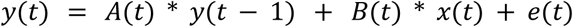

where * represents a convolution, and e(t) is an additive innovation process, or equivalent to an uncorrelated error in the linear prediction. A first-order VARX model (with filter length n_a_=n_b_=1) is equivalent to the classic first-order linear dynamical system model used extensively in the neuroscience literature, e.g. ^38,39^. When no exogenous input is modeled (n_b_=0), we are left with endogenous recurrent dynamic alone, which is the same as the vector autoregressive models (VAR) used in the context of “Granger causality”, e.g. ^41,87^. In the z-domain or the frequency domain this can be written as H(ω)=(1-A(ω))^−1^B(ω) -- this motivated us here to show the effect of the recurrent component separately, (1-A)^−1^ , in Fig. 2c-d, to visualize the endogenous dynamic. Figure 4.c-g then compares H with B.

The estimation of the filter parameters A(t) and B(t) follows ordinary least squares, i.e. minimize the mean squared error. The filter matrix A(t) and B(t) have dimensions [n_a_,d_y_,d_y_] and [n_b_,d_y_,d_x_], respectively, where n_a_ and n_b_ are the number of tabs in each filter, d_y_ is the dimensions of the dynamical endogenous signals y(t) and d_x_ is the dimension of the exogenous inputs signals x(t). Testing for statistical significance of each of the d_y_*d_y_ and d_y_*d_x_ filters in matrices A and B are based on the Granger formalism. This formalism provides effect size matrices, generalized R_A_ and R_B_, respectively, which measure how well a signal source (predictor “cause”) can linearly predict another signal (predicted “effect”) from its history. Therefore, any pair of signals has two directions of prediction. Effect sizes for all signals therefore form a matrix of effect sizes (Fig. 2). When the effect size is statistically significant, we will refer to this as a “connection”. In the case of R_A_, the effect size matrix is square but not necessarily symmetric, reflecting asymmetric directed connectivity. Please note that the choice of words of “cause”, “effect” and “connection” are purely for convenience and we do not mean to imply an actual physical connection that mediates a cause and effect relationship. We are not making any claims on causality, because of the assumption of second-order stationarity of signals required for this statistical test. The signals here almost certainly are not stationary in their power. For a more thorough discussion of this important limitation, please see ^37^.

#### Significance testing for VARX parameters (A and B filters)

Statistical significance of the endogenous (A) and exogenous (B) filters was assessed using the Granger test, following the implementation ^36,37^.

For each predictor-effect pair, the model compares 1. Full model error σ_f_ where all predictors are included, 2. Reduced model error σ_r_, where the predictor is removed. This yields the deviance

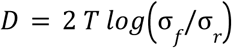

Which is Χ^2^-distributed for large T. The analytical p-value is

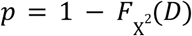

Effect size is given by the generalized R^2^

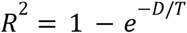

Thus, each filter of matrix A and B yields a positive effect-size matrix R_A_ and R_B_. In figures 2b, 4b, 4e-g we only display a connection when p < 0.001 and R>0.02 for better visualization.

#### Group-level inference

Models were estimated separately for each participant, using identical regularization (λ = 0.01), filter lengths (n_a_, n_b_), and preprocessing steps. Group summaries were computed by averaging R_A_ or R_B_ across subjects. Group significance is therefore based on the consistency of signed effect sizes and the distribution of p-values across subjects, consistent with the approach in ^36^ (see Methods pp. 15-16 and Appendix 3)

#### Statistical comparison of effect-size matrices

To quantify whether two directed effect-size matrices shared a similar interaction pattern, we used a row/column-permutation similarity test based on the Mantel test for similarity ^88^. The Mantel test compares the similarity of two matrices *X* and *Y*, with a null distribution obtained by the similarity of *X* and randomly row/column-shuffled versions of *Y*. As such, it rules out whether the observed similarly between *X* and *Y* is due to the chance occurrence of the alignment of the elements of *Y* with respect to *X*.

Given that the standard Mantel test was designed to test the similarity of two matrices containing pairwise distances of a number of objects, here we use a modified version of the Mantel test, while keeping its operating principle intact. First, we normalize each matrix by its Frobenius norm, to compare the connectivity patterns regardless of scale. Second, we use a symmetrized Tanimoto similarity metric as follows. The usual Tanimoto similarity of two vectors *x* and *y* is defined as 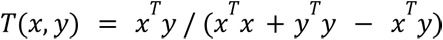. When applied to two matrices *X* and *Y*, we interpret *T*(*X*, *Y*) as the Tanimoto similarity between vectorized versions of *X* and *Y* ^89^. Let π and π_2_ be two random permutations of *N* objects and let *X*^π1^ and *Y*^π2^ denote the row/column-permuted versions of *X* and *Y* via π_1_ and π_2_, respectively. Then, we define the symmetrized Tanimoto similarity as 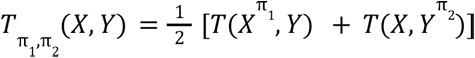. Letting *I* denote the identity permutation, we compare *T_I,I_*(*X*, *Y*) against a null distribution obtained from 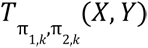 by drawing *K* random permutation pairs (π_1*,k*_, π_2,*k*_), *k* = 1, 2, ···, *K*. The null distribution from this shuffle is shown in Fig. S8a-c (red line is the identity permutation). The test p-value is then computed as 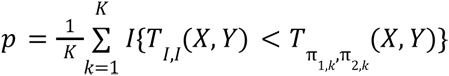. Code for this was added to the VARX toolbox.^90^

#### Directional asymmetry analysis

To quantify directional asymmetry in brain–body interactions, we fit the VARX model independently for each participant, yielding subject-specific directional coupling matrices *R_A_*.

This approach treats participants as independent samples and captures between-subject variability in estimated interactions. For each participant *s*, we identified paired bidirectional connections linking the same brain and peripheral signals and computed a scalar asymmetry measure defined as the mean paired difference in mean effect size between brain→body and body→brain directions:

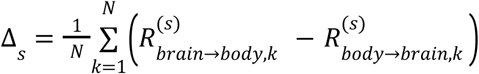

where *k* indexes corresponding directional connections and *N* is the total number of paired connections. This yielded one asymmetry estimate per participant. Statistical significance was assessed across participants using a Wilcoxon signed-rank test (one-sided), testing the null hypothesis that paired brain→body and body→brain coupling strengths do not differ. To rule out that the asymmetry reported in the results section is not due to the differing spectra in these signals (i.e. spike trains are difficult to linearly regenerate from slow signals), we repeated this analysis after low-pass filtering the spike trains coding saccade onsets (by convolving with a 300 ms-wide Gaussian kernel to match the peak power of the EEG), and high-pass filtering all the behavioral and physiological signals with the same 0. 3 Hz high-pass filter that was applied to the EEG. The statistical results are largely unchanged.

#### Model order estimation

To determine the length of filter *n_a_* and *n_b_* , we used here the Akaike Information Criterion (AIC) ^91^ for each channel in *y*(*t*):

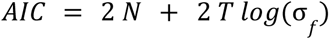

Where, *N* = *n_a_ d_y_* + *n_a_ d_y_*, is the number of model parameters for each channel and 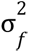 is the estimation error variance. Note that as *N* grows, i.e., considering longer filters, the first term in *AIC* increases, but the longer filters improve estimation and thus 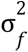 and the second term of *AIC* decrease. As such, minimizing *AIC* accounts for a balance between model complexity (i.e., longer filters) and estimation performance (i.e., lower variance).

A single set of parameters n_a_ and n_b_ can be selected by taking the sum of AIC over all d_y_ channels. Code for this was added to the VARX toolbox.^90^ We used the data during the attentive listening condition for Experiments C and F independently when selecting optimal parameters (Fig. S6). For *n_b_*, we picked the value with the lowest AIC. At a sampling frequency of *f_*s*_* = 25Hz, we found *n_b_* = 39 and 26 for Experiments C and F. However, for *n_a_* the AIC did not show a unique minimum. Instead, increasing *n_a_* produced a nearly linear decrease in AIC without reaching an identifiable optimum. This behavior likely reflects nonlinear long-term memory effects in the data, which linear autoregressive models are not suited to capture.

To avoid overfitting while still modeling short-timescale dependencies, we selected *n_a_* = 10, corresponding to the knee point at which AIC transitions into its approximately linear regime (Fig. S6). This choice produced stable results across experiments and sampling rates.

We further evaluated sensitivity to sampling frequency by fitting models at 10 Hz and 25 Hz. Downsampling to 10 Hz removed alpha-band activity (∼10 Hz), which substantially reduced effect sizes *R*_A_ between EEG electrodes, while leaving the variance explained by other electrodes largely unchanged. This indicates that low-frequency EEG components drive most predictive power, whereas spatial coupling patterns in *R_A_* depend partly on oscillatory structure such as alpha coherence.

#### Missing values

Missing values were excluded from the cross-subject statistical analyses. Missing samples and artefacts in the data were in general removed and replaced with interpolated values, the details of which are discussed in each section above. While the VARX estimation of linear prediction parameters excluded samples with NaN anywhere in the prediction window.

## Supplemental Information

**Figure S1.**
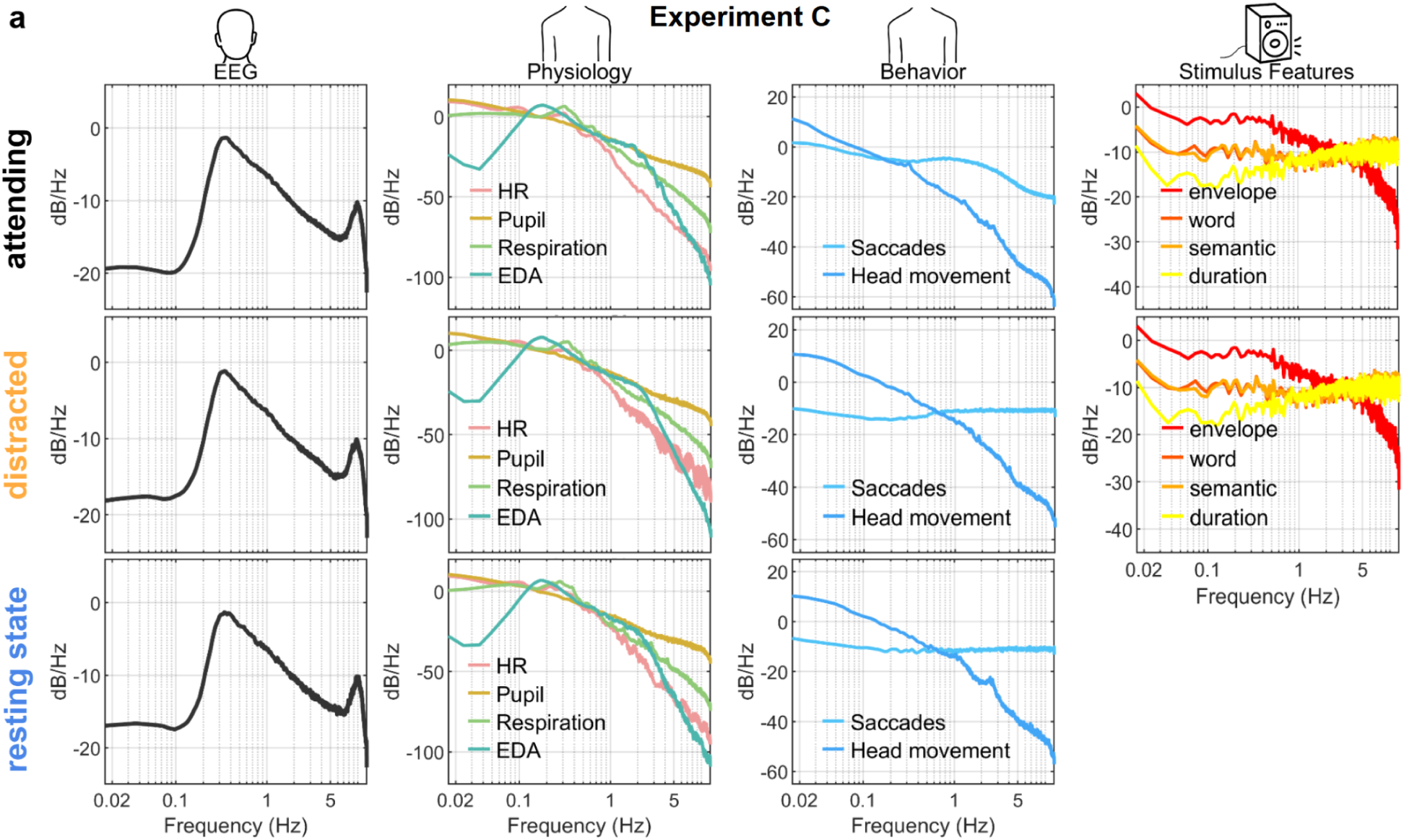
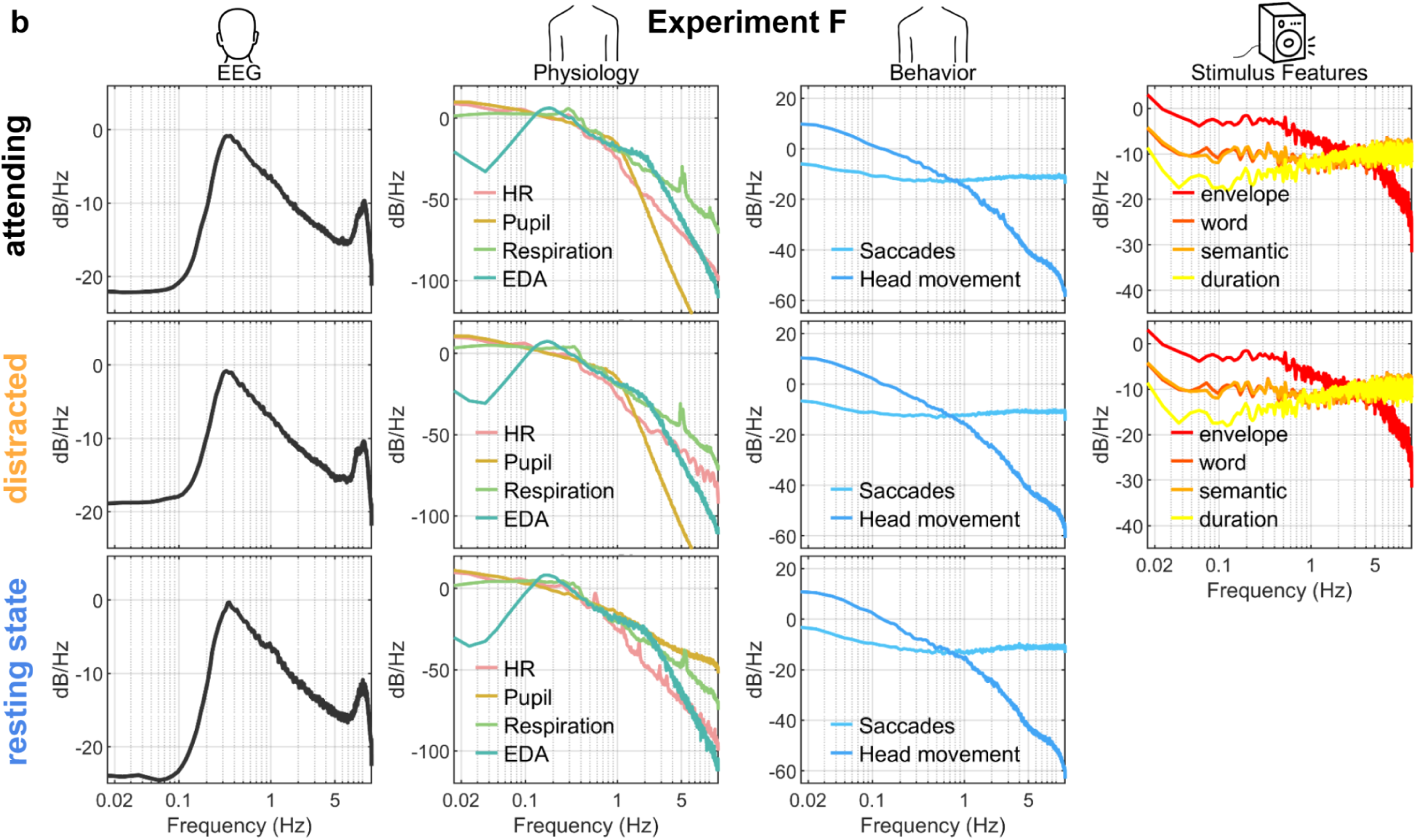
Power spectra of neural, physiological, behavioral, and stimulus signals across conditions and experiments. This figure complements Figure 1 by showing power spectra for all recorded neural, physiological, behavioral, and stimulus signals separately for attentive listening, distracted listening, and resting state, and for both Experiment C (constrained gaze) and Experiment F (free gaze). EEG slow potentials show dominant power at infra-slow frequencies across all conditions. Physiological signals (heart rate, pupil diameter, respiration, electrodermal activity) and behavioral signals (saccades, head motion) likewise exhibit substantial low-frequency structure. Speech features span a broad frequency range, with the acoustic envelope showing broadband fluctuations and linguistic features exhibiting both slow and faster components. Power spectral profiles are highly consistent across cognitive states and experiments, demonstrating that differences reported in the main text are not driven by gross differences in signal power or frequency content.

**Figure S2.**
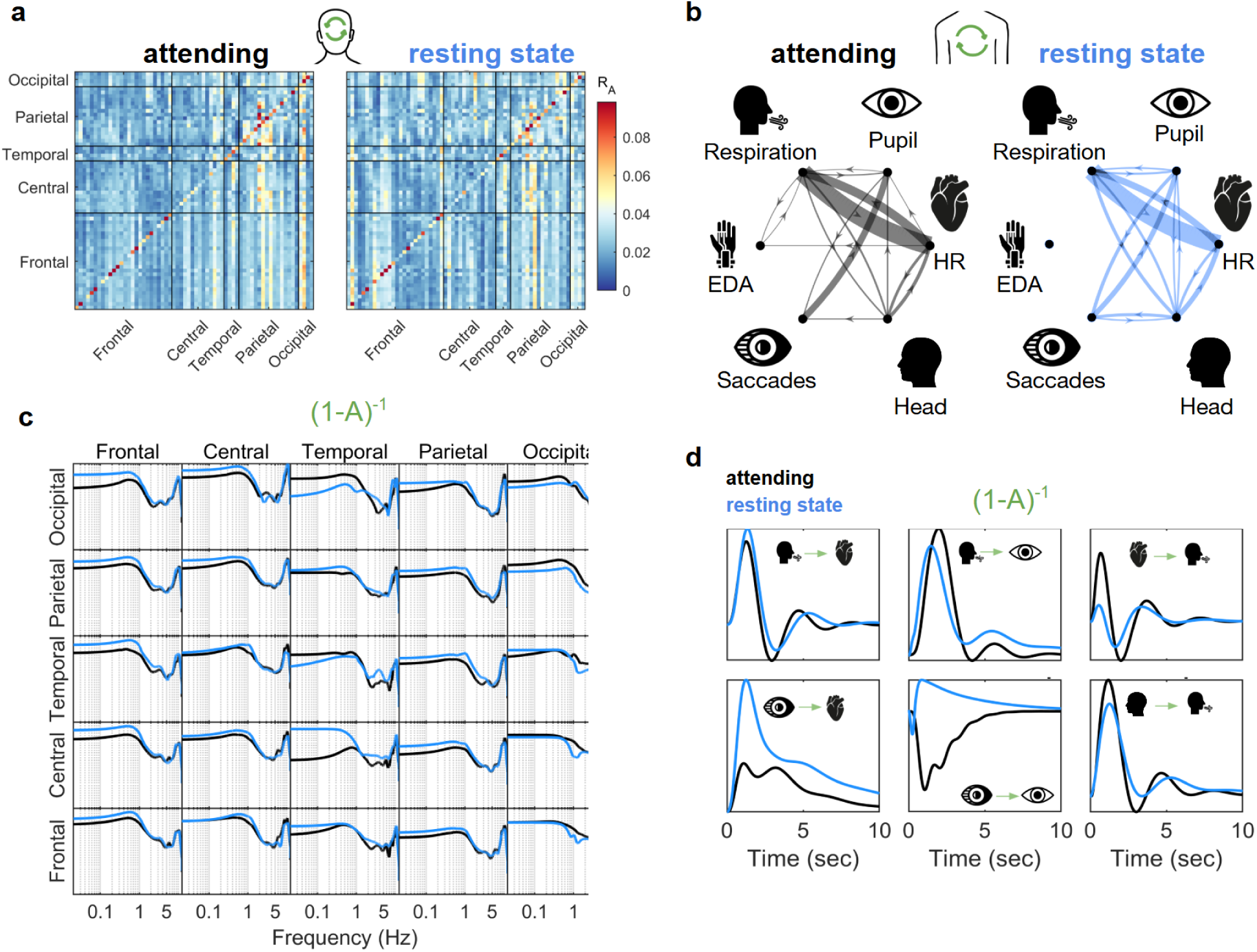
Recurrent brain-body dynamics replicate across experiments. This figure reproduces the analyses shown in Figure 2 for Experiment F (free gaze), demonstrating that the stability of endogenous brain-body dynamics generalizes across experimental setups. **(a)** Intrinsic interaction matrices A estimated from the VARX model for EEG signals during attentive narrative listening (left) and resting state (right). As in Experiment C, the spatial structure of recurrent EEG interactions across frontal, central, temporal, parietal, and occipital regions is highly similar between listening and rest, indicating preserved endogenous cortical dynamics. (b) Intrinsic interactions among peripheral physiological and behavioral signals during attentive listening (left) and resting state (right). Directed edges indicate significant endogenous influences between respiration, pupil diameter, heart rate (HR), electrodermal activity (EDA), saccades, and head motion, with edge thickness proportional to effect size (R_A_). Network structure closely matches that observed in Experiment C. **(c)** Frequency-domain magnitude responses of recurrent EEG interactions, averaged over electrode pairs within each cortical region, corresponding to the recurrent component (1−A)^−1^ (impulse responses shown in Fig. S7b). **(d)** Time-domain impulse responses between peripheral physiological and behavioral signals derived from (1−A)^−1^, shown for attentive listening (black) and resting state (blue). As in Experiment C, impulse response shapes and timescales are largely preserved across cognitive states.

**Figure S3.**
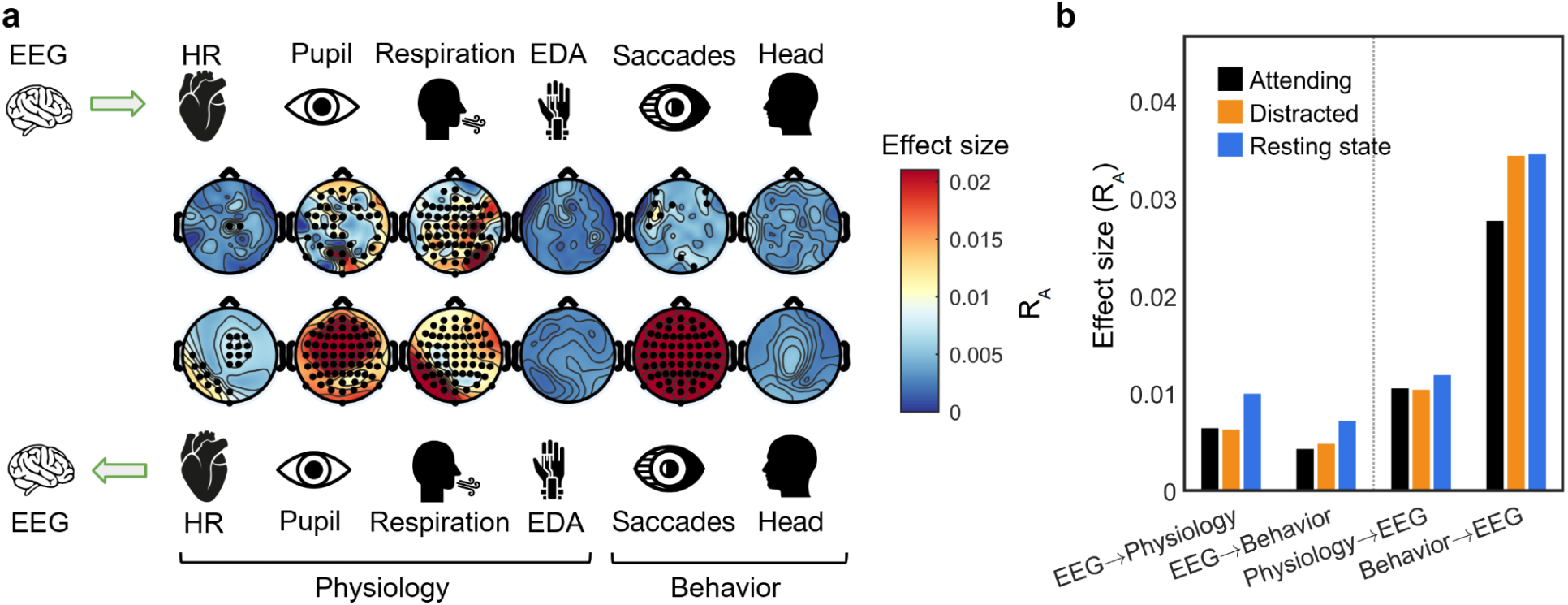
Directional asymmetry between brain and body replicates across experiments. This figure reproduces the analyses shown in Figure 3 for Experiment F (free gaze). **(a)** Topographic maps show the effect size (R_A_) of intrinsic interactions between EEG and peripheral physiological and behavioral signals during attentive listening. The top row depicts influences from EEG to heart rate (HR), pupil diameter, respiration, electrodermal activity (EDA), saccades, and head motion. The bottom row shows the reverse direction, from each peripheral or behavioral signal to EEG. Black dots indicate electrodes with statistically significant effects. As in Experiment C, influences from body and behavior to EEG are broader and stronger than those in the opposite direction. **(b)** Grouped bar plots summarize directed effect sizes between EEG and peripheral physiology or behavior across conditions (attentive, distracted, resting state). Values are averaged across EEG electrodes and channels within each category. Consistent with the main experiment, body→EEG and behavior→EEG effects exceed EEG→body effects across cognitive states.

**Figure S4.**
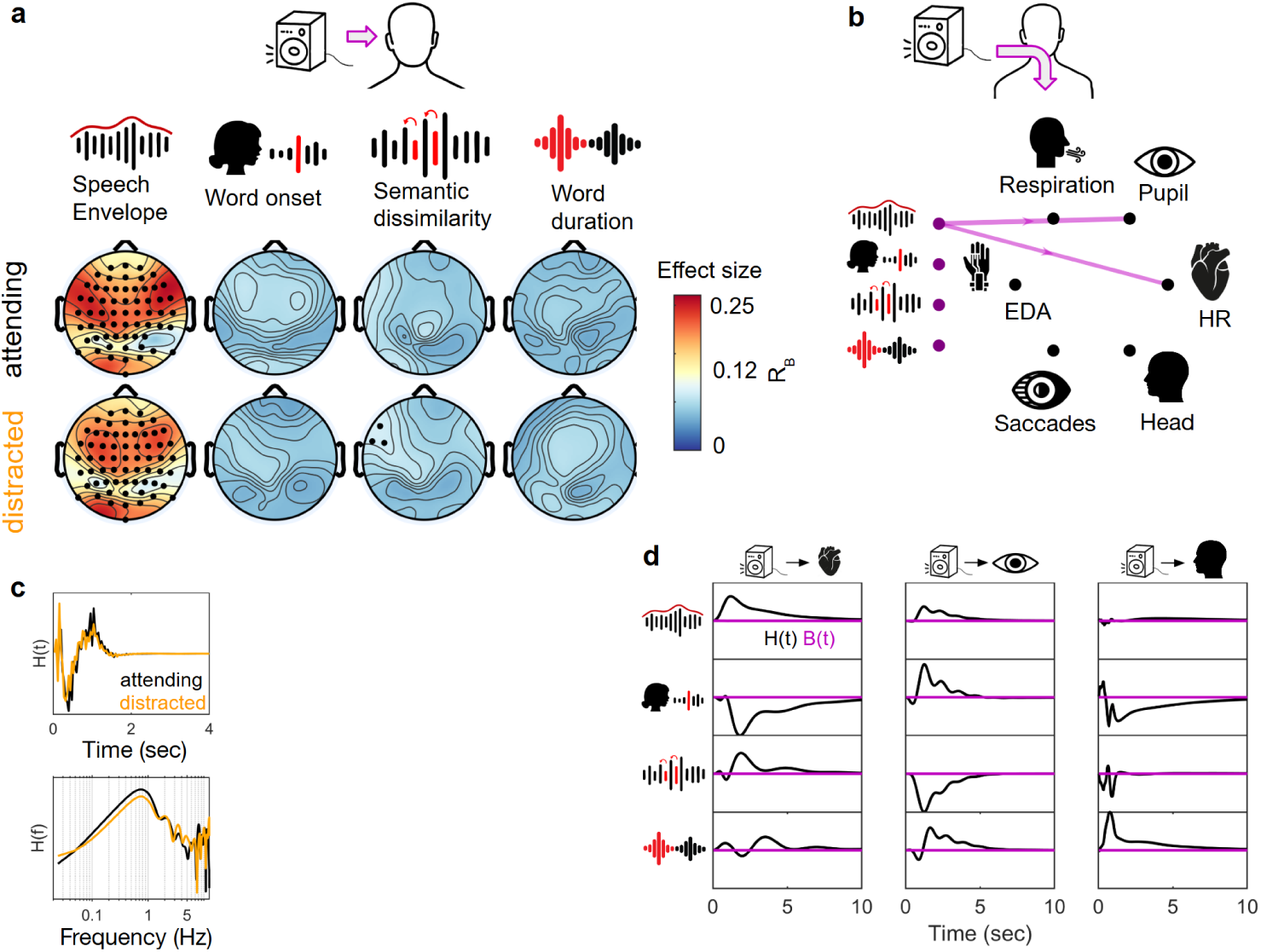
Direct and indirect stimulus-driven effects on brain–body dynamics replicate across experiments. This figure reproduces the analyses shown in Figure 4 for Experiment F (free gaze). **(a)** Topographic maps show the effect size (R_B_) of stimulus-driven responses from the VARX extrinsic filter B(t) for four speech features: speech envelope, word onsets, semantic dissimilarity, and word duration. Maps are shown for attentive (top row) and distracted (bottom row) listening. Black dots indicate electrodes with statistically significant extrinsic effects. Compared to Experiment C, fewer electrodes reached significance for higher-level linguistic features, while robust envelope-driven responses were preserved. **(b)** Network summary of significant extrinsic stimulus-to-physiology effects. Edges indicate significant direct pathways from speech features to physiological signals, with line thickness proportional to effect size. As in the main experiment, direct autonomic effects are largely restricted to low-level acoustic fluctuations. **(c)** Time-domain impulse responses H(t) for selected EEG electrodes with significant stimulus-driven effects, plotted separately for attentive (black) and distracted (orange) listening. **(d)** Corresponding frequency-domain representations H(f), illustrating similar spectral structure across conditions despite reduced statistical power. **(e-g)** Time-domain impulse responses for significant stimulus-to-physiology effects, showing the direct component B(t) (magenta) and the full system response H(t) (black). As in Experiment C, physiological responses primarily emerge through reverberation within the endogenous dynamics rather than direct stimulus drive.

**Figure S5.**
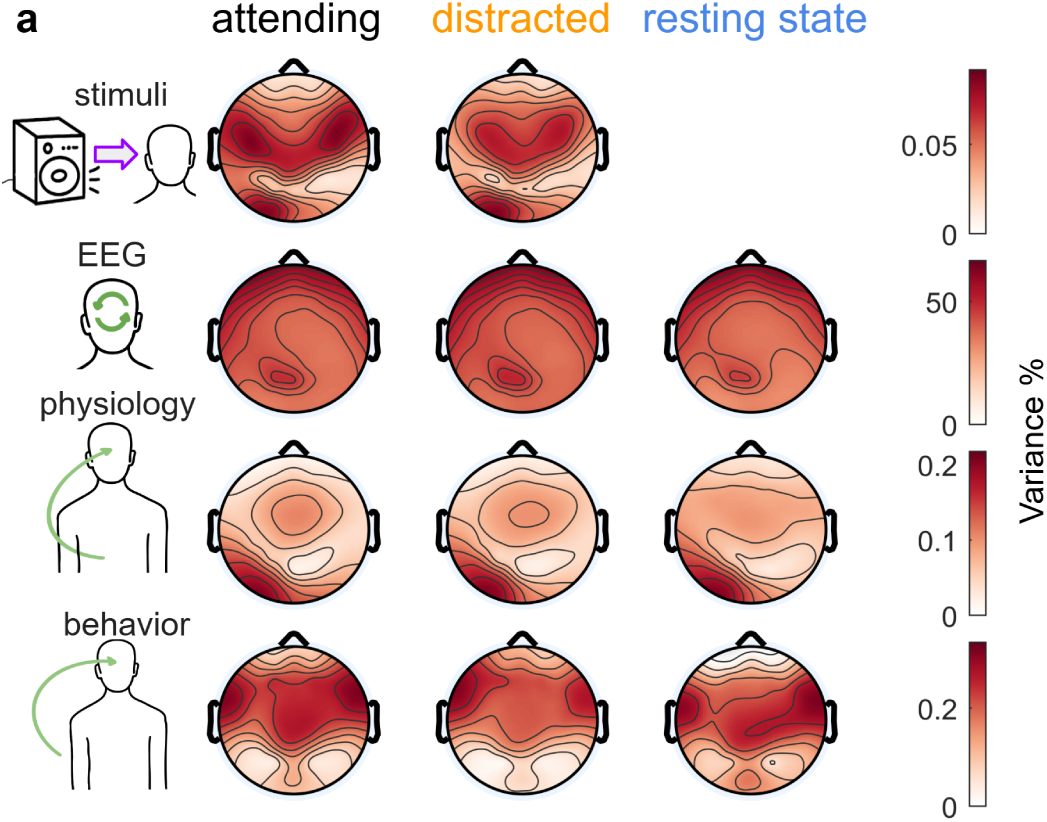
EEG variance decomposition replicates across experiments. This figure reproduces the EEG variance decomposition shown in Figure 5 for Experiment F (free gaze). Topographic maps show the percentage of EEG variance explained by four components of the VARX model: external stimulus features (top row), intrinsic EEG dynamics (second row), intrinsic influences from peripheral physiology (third row), and behavior (bottom row). Maps are shown for attentive listening, distracted listening, and resting state. As in Experiment C, intrinsic EEG dynamics account for the largest fraction of variance across the scalp, with smaller but reliable contributions from behavioral and physiological signals and minimal variance explained directly by the stimulus. The spatial distribution and relative magnitudes of these components are highly consistent across experiments, despite differences in gaze constraints.

**Figure S6.**
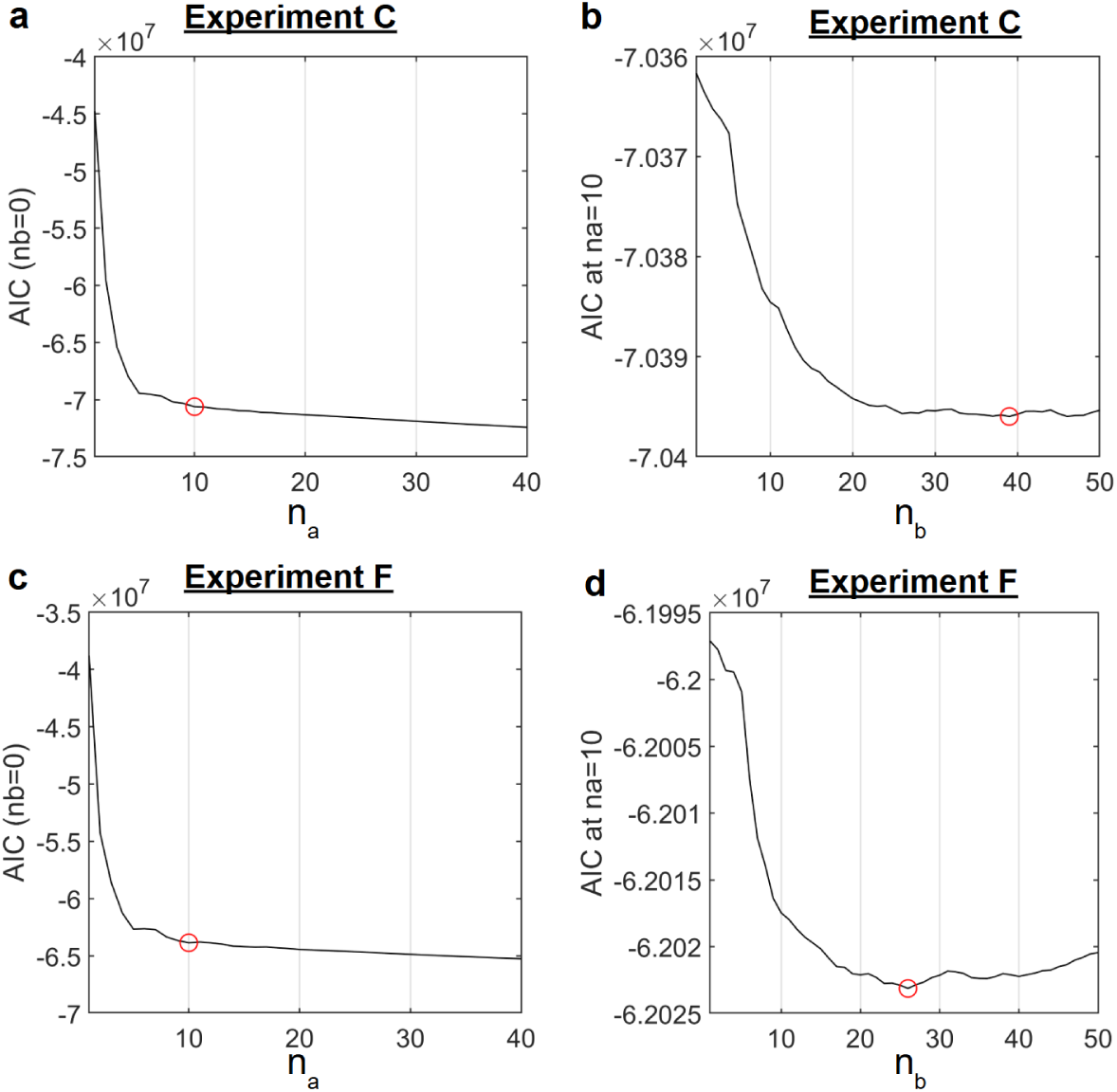
Akaike Information Criterion (AIC) for model selection. AIC was calculated, and optimal values for model order n_a_ and n_b_ were selected on the attentive listening task, separately for experiments C and F (to test reproducibility). First, n_a_ was fit with no input, i.e., a VAR model, then, with the optimal n_a_ VARX was fit to find the best n_b_. Optimal values are indicated with a red circle. The choice of n_a_=10 was based on the knee point of the AIC in Experiment C, in lieu of an actual minimum value. Experiment F gave the same knee point.

**Figure S7.**
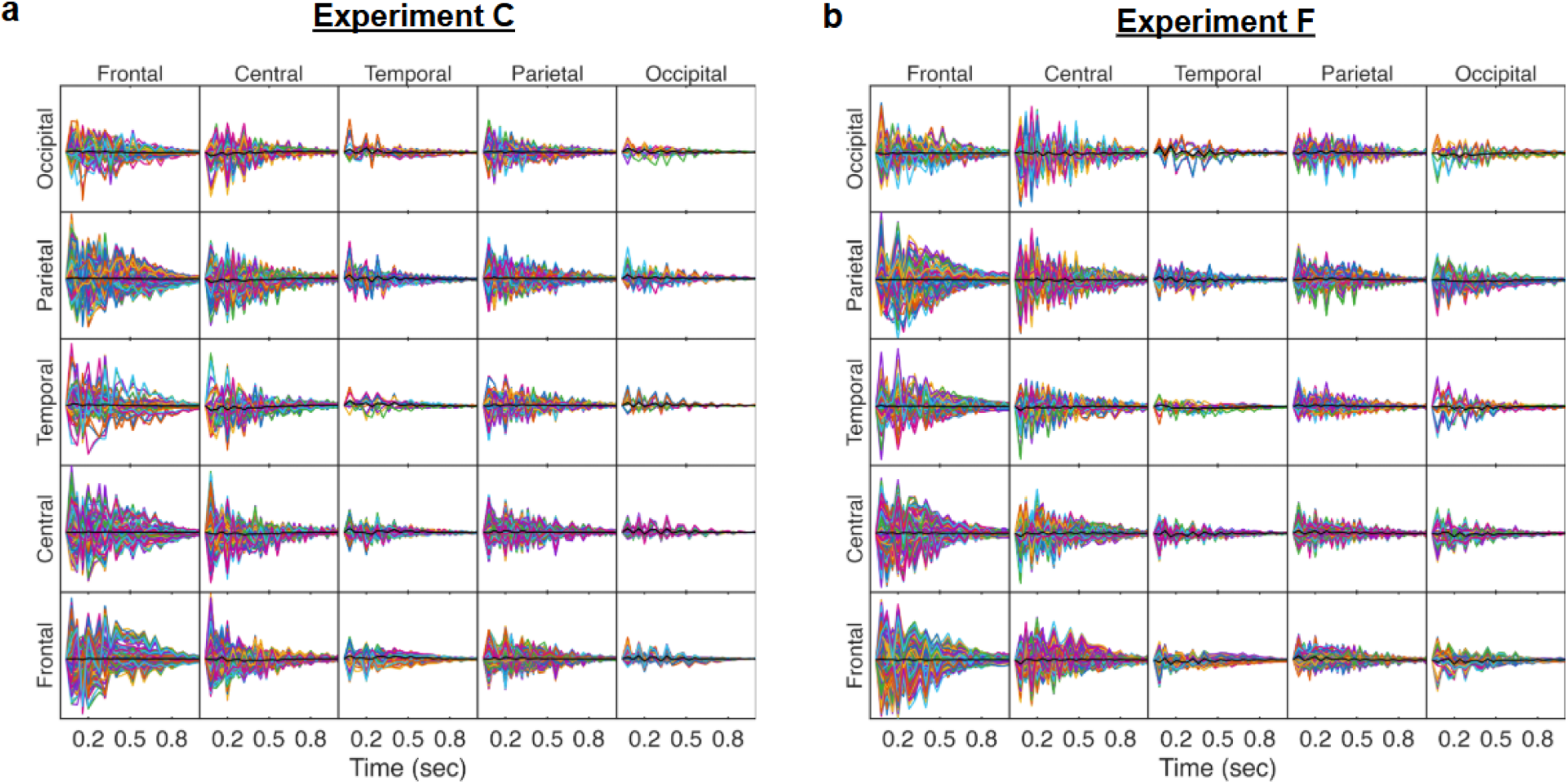
Impulse responses between EEG electrodes. Each curve is the impulse response from one electrode to another (columns are the predictor, and rows are the predicted, i.e. input and output of the impulse response). The black line shows the average across electrodes in that region pair. The impulse responses are not phase aligned. Impulse responses here are computed with VARX model parameters (1-A)^−1^, fit to the attending condition.

**Figure S8.**
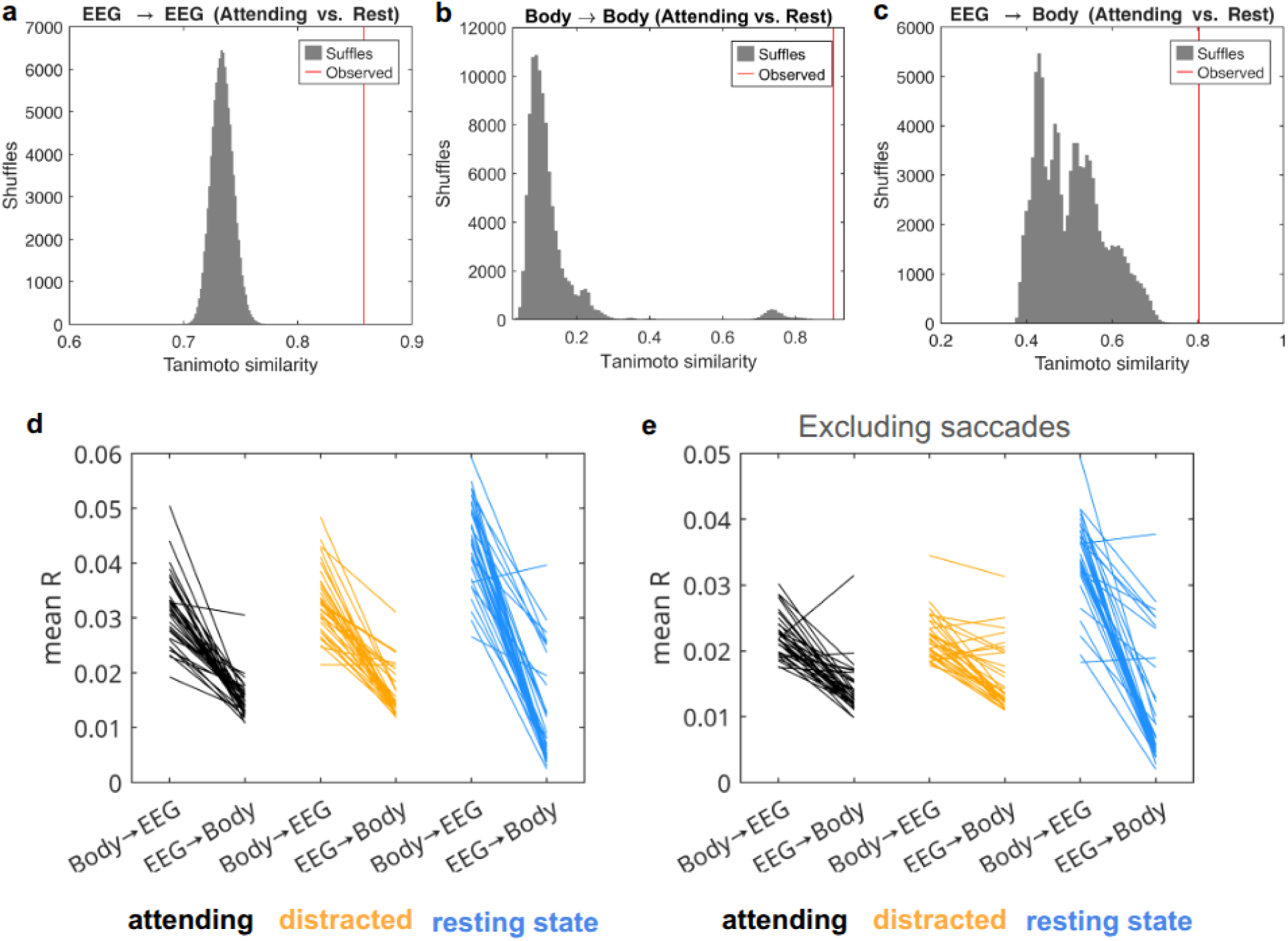
Permutation tests for stability of interaction structure across conditions and directional asymmetry of brain-body coupling. **(a-c)** Row/column relabeling permutation test comparing the structure of directed effect size matrices R_A_ between attentive listening and rest. For each comparison, matrices were Frobenius-normalized (entries outside the analysis mask set to zero; NaNs set to zero) and similarity was quantified with a symmetrized Tanimoto similarity. The null distribution (gray histogram, “Shuffles”) was generated by independently permuting row and column labels (random relabeling of sources and targets). The red line shows the observed similarity. The null distributions are non-Gaussian, supporting the need for shuffling procedure; the null distribution sometimes centers around 0.2 and sometimes around 0.75, so it adapts to the actual matrices being tested and thus we are testing against a relevant null distribution, not a generic one. Panels show comparisons for EEG→EEG (a), Body→Body (b), and EEG→Body Small p-values indicate the matrices are more similar than expected under random relabeling. **(d)** Mean effect size R for prediction of EEG from physiological and behavioral signals (Body → EEG) and the reverse EEG→Body. Each line is the result of a VARX model fitted independently to each subject (here Experiment C, N=38). Wilcoxon rank sign test for the three conditions: Z =5.23, 5.09, 5.29, p = 1.68·10^−7^, 3.65·10^−7^, 1.24·10^−7^. **(e)** Same as d, but excluding the saccades from the mean R calculation (but not the model fit): z = 4.89, 4.80, 5.26, p = 1.03·10^−6^ , 1.6·10^−6^, 1.5·10^−7^

